# Persistent Interruption in Parvalbumin Positive Inhibitory Interneurons: Biophysical and Mathematical Mechanisms

**DOI:** 10.1101/2024.03.04.583352

**Authors:** Carol M Upchurch, Christopher J Knowlton, Simon Chamberland, Carmen C Canavier

## Abstract

Persistent activity in principal cells is a putative mechanism for maintaining memory traces during working memory. We recently demonstrated persistent interruption of firing in fast-spiking parvalbumin-expressing interneurons (PV-INs), a phenomenon which could serve as a substrate for persistent activity in principal cells through disinhibition lasting hundreds of milliseconds. Here, we find that hippocampal CA1 PV-INs exhibit type 2 excitability, like striatal and neocortical PV-INs. Modelling and mathematical analysis showed that the slowly inactivating potassium current K_v_1 contributes to type 2 excitability, enables the multiple firing regimes observed experimentally in PV-INs, and provides a mechanism for robust persistent interruption of firing. Using a fast/slow separation of times scales approach with the K_v_1 inactivation variable as a bifurcation parameter shows that the initial inhibitory stimulus stops repetitive firing by moving the membrane potential trajectory onto a co-existing stable fixed point corresponding to a non-spiking quiescent state. As K_v_1 inactivation decays, the trajectory follows the branch of stable fixed points until it crosses a subcritical Hopf bifurcation then spirals out into repetitive firing. In a model describing entorhinal cortical PV-INs without K_v_1, interruption of firing could be achieved by taking advantage of the bistability inherent in type 2 excitability based on a subcritical Hopf bifurcation, but the interruption was not robust to noise. Persistent interruption of firing is therefore broadly applicable to PV-INs in different brain regions but is only made robust to noise in the presence of a slow variable.

**Significance Statement:** Persistent activity in neuronal networks is thought to provide a substrate for multiple forms of memory. The architecture of neuronal networks across many brain regions involves a small number of inhibitory neurons that control many principal neurons. We propose that persistent silencing of fast-spiking parvalbumin-expressing inhibitory interneurons (PVINs) can result in persistent activity of principal neurons. We use a mathematical approach and computer simulations to investigate the mechanisms governing persistent interruption of firing in hippocampal and cortical PV-INs. We show how a slowly changing state of a particular ion channel controls the long-lasting silence imposed by persistent interruption. Overall, our results provide a conceptual framework that positions the persistent interruption of PV-INs firing as a potential mechanism for persistent activity in principal cells.

## Introduction

The hippocampus has been implicated in short-term memory tasks (Leavitt et al. 2017) in which information retention is necessary for a few hundreds of milliseconds (Weiss et al. 1999) to tens of seconds (Hampson et al. 2004; Hampson and Deadwyler 2000) in the absence of sensory stimulation. For example, cue associated activity persists during the delay period of delayed response tasks in multiple regions of the brain (Leavitt et al. 2017), including the hippocampus (Cahusac et al. 1989; Watanabe and Niki 1985). A possible substrate for the representation of information in the absence of a sensory stimulus is persistent activity, defined as “a sustained change in action potential discharge that long outlasts a stimulus” (Major and Tank 2004). Multiple mechanisms at both the network and at the single cell level have been proposed to underly persistent neuronal activity. One proposed mechanism for persistent activity is a mnemonic attractor sustained by reverberatory dynamics through feedback loops in a neural assembly (Wang 2021). Another putative substrate for persistent activity relies on the intrinsic bistable dynamic of individual neurons (Zylberberg and Strowbridge 2017). On the single cell level, persistent Na^+^ current (Yamada-Hanff and Bean 2013) in CA1 pyramidal cells, Ca^2+^ activated Ca^2+^ currents in entorhinal cortical pyramidal cells (Egorov et al. 2002) and CA1 pyramidal cells (Combe et al. 2023), and low threshold Ca^2+^ currents in other cells (Otsuka et al. 2001; Perrier et al. 2002) have been shown to contribute to regenerative firing, by making the neuron bistable or even multi-stable, and thus more likely to continue firing after excitation is terminated.

Neuronal discharge was originally described in terms of firing continuity as a function of continuous current injection. Neurons with Hodgkin’s type 1 excitability can fire arbitrarily slowly and have a continuous frequency/current relationship. In contrast, neurons with Hodgkin’s type 2 excitability show repetitive firing of action potentials that cannot be sustained below a threshold firing frequency, and the frequency/current relationship is discontinuous at the value of applied current that is sufficient to sustain repetitive firing (Hodgkin 1948). The determinant of the excitability type is whether inward or outward currents are dominant at equilibrium at the values of membrane potential traversed during the interspike interval. In type 1, inward currents dominate, whereas in type 2 outward currents dominate at equilibrium. Therefore, the interspike interval must be short enough that the slower outward currents do not come to equilibrium, resulting in a lower bound on the frequency that can be sustained during repetitive firing. Neurons with type 2 excitability display inherent bistability near the abrupt onset of tonic firing and could contribute to switch-like transitions in neuronal networks, but whether and how it can sustain persistent activity remains understudied.

We suggest a mechanism for persistent activity in pyramidal cells (PYR) involving the persistent silencing of presynaptic interneurons in a simple feedforward inhibitory circuit. Recent work has shown that a GABA_A_-mediated inhibitory post-synaptic potential can interrupt tonic firing in hippocampal parvalbumim positive interneurons (PV-INs) for hundreds of milliseconds after the IPSP has dissipated (Chamberland et al. 2023b), a phenomenon supported by an interplay between the slowly-decaying IPSP and rebalancing of membrane conductance including K_V_1. The resulting long-lasting silence in hippocampal PV-INs tripled the firing rate of postsynaptic CA1-PYRs. Therefore, the mechanisms controlling firing interruption in hippocampal PV-INs could be key in understanding persistent activity in downstream CA1-PYRs.

Here, we used single-compartment conductance-based computational models of PV-INs to describe the persistent interruption of firing dynamics. The model replicated type 2 excitability and explained the different firing regimes experimentally observed in CA1 hippocampal PV-INs. Bifurcation analysis and a fast/slow separation of time scales approach (Bertram and Rubin 2017) revealed a subcritical Hopf bifurcation dependent on the slow dynamics of K_V_1 inactivation. While K_v_1 was not necessary for firing interruption in a mEC PV-IN model, it conferred significant robustness to noise.

## Methods

### Acute hippocampal slice preparation and electrophysiological recordings

All experiments performed were approved by NYU Langone IACUC protocols. Acute brain slices were prepared from P20 – P35 male and female mice. *Pv*-Ai9 animals were obtained by crossing Pv-Cre mice (B6;129P2-Pvalb^tm1(cre)Arbr^/J, JAX stock #017320) to Ai9 reporter line mice (B6.Cg-Gt(ROSA)26Sor^tm9(CAG-tdTomato)Hze^/J, JAX stock #007909). Mice were deeply anesthesized with isoflurane and decapitated. The brain was extracted and placed in ice-cold oxygenated (95% O2 and 5% CO2) sucrose ACSF solution. The sucrose ACSF solution contained (in mM): sucrose 185, NaHCO_3_ 25, KCl 2.5, NaH2PO4 1.25, MgCl_2_ 10, CaCl_2_ 0.5, and glucose 25 (pH = 7.4, 330 mOsm). The brain was dissected and transverse acute hippocampal slices (300 μm) were prepared on a vibratome (Leica VT1000S). Hippocampal slices were transferred to an ACSF-containing recovery chamber that was continuously oxygenated and maintained at 32°C. The ACSF contained (in mM): NaCl 125, NaHCO_3_ 25, KCl 2.5, MgCl_2_ 2, CaCl_2_ 2, and glucose 10 (pH = 7.4, 300 mOsm). After a recovery period of 30 minutes, the temperature was allowed to come down to room temperature for the rest of the experiments. Experiments were started no less than one hour after slice transfer to the recovery chamber. For electrophysiological recordings, acute slices were transferred to a submerged recording chamber under an upright microscope (BX61WI, Olympus) equipped with a water-immersion 40X objective (Zeiss). TdTomato+ interneurons located in the CA1 strata oriens bordering the pyramidal cell layer were visually identified and selected for whole-cell recordings. Recording electrodes were prepared from borosilicate glass filaments using a micropipette puller (Sutter). Recording electrodes were filled with a K+-based intracellular solution that contained (in mM): 130 K-gluconate, 10 HEPES, 2 MgCl_2_.6H_2_O, 2 Mg_2_ATP, 0.3 NaGTP, 7 Na_2_-Phosphocreatine, 0.6 EGTA, 5 KCl. The pH was adjusted to 7.2 using 1 M KOH, and the final solution had an osmolality of 295 mOsm. Recording electrodes had resistance of 3 – 6 MΩ. The electrophysiological signal was amplified with an Axopatch 200B and digitized at 10 kHz with a Digidata 1322A (Axon Instruments). Current-clamp recordings consisted of a 500 ms depolarizing current step of increasing amplitude, delivered over multiple trials every 10 seconds. The data was recorded on a personal computer.

### Analysis of electrophysiological data

The input resistance was measured using a voltage step from -60 mV to -50 mV. The time constant was found using a current step yielding approximately a 10mV change in membrane potential and assuming a mono-exponential fit. The resulting average capacitance was calculated using the input resistance and the time constant: τ=RC.

### Computational Model

We began with the PV+ FS model from (Chamberland et al. 2023b). The differential equation for the membrane potential (V in mV) for the neuron is given by: *C*_*M*_*dV*/*dt* = −*I*_*App*_ − *I*_*leak*_ − *I*_*Kdr*_ − *I*_*Kv*1_ − *I*_*Na*_ where C_M_ is the capacitance of the cell membrane (1µF/cm^2^), *I_App_* is the current injected, *I_leak_* is the passive leak current, *I_K_V_3_* is the fast delayed rectifier, *IK_v_1* is the slowly inactivating current mediated by K_v_1, and *I_Na_* is the fast sodium current. *I_leak_* was described by the equation *I*_*leak*_(*v*) = *Ig*_*L*_ * (*v* − *E*_*L*_). The differential equations for the gating variables are of the form dx/dt = (x-x_inf_(*v*))//τ_x_(*v*). *I_Kv3_* was described by the following equations: *I*_*Kv*3_(*v*, *n*) = *g*_*Kv*3_ * *n*^2^ * (*v* − *E*_*K*_) where *n* relaxes to its instantaneous voltage state, *n*_*inf*_(*v*) = {1 + exp [−(*v* + 12.4)/6.8]}^−1^ with time constant *r*_*n*_(*v*) = {0.087 + 11.4 * {1 + exp [(*v* + 14.6)/8.6]}^−1^} * {0.087 + 11.4 * {1 + exp [−(*v* − 1.3)/18.7]}^−1^} and *g_Kv3_* retained its original value of 0.223 mS/cm^2^. Kv3 channels are critical to models of fast spiking interneurons because they deactivate very quickly enabling very fast spiking (Rudy et al. 1999; Rudy and McBain 2001), and are present in fast spiking PV+ inhibitory interneurons at a much higher level than in other cell types (Erisir et al. 1999). Their contribution to the AHP also limits the accumulation of Na^+^ channel inactivation (Erisir et al. 1999).

The sodium current was described by *I*_*Na*_(*v*, *m*, *h*) = *g*_*Na*_ * *m*^3^ * *h* * (*V* − *E*_*Na*_) where *E_Na_*= 50 mV, *m*_*inf*_(*v*) = {1 + exp [−(*v* + 24)/11.5]}^−1^, τm= 0.001ms, *h*_*inf*_(*v*) = {1 + exp [(*v* + 58.3)/6.7]}^−1^, *r*_*h*_(*v*) = 7 * {1 + exp [(v + 60)/12] }^−1^ and *g_Na_* retained its original value 0f 0.1125 mS/cm^2^. The half activation for the activation gate of the sodium channel was changed from -24mV to -22mV to more resemble the rheobase of PV+ interneurons in CA1. *I*K_v_1 was described by the equation *I*_*Kv*1_ = *g*_*Kv*1_ * *p* * *q* * (*V* − *E_K_*) where E_K_ = -90 mV. The activation gate *p* was described by *p*_*inf*_(*v*) = {1 + exp [−(*v* + 41.4)/26.6]}^−4^ with *r*_*p*_ = 0.448 *ms*. The inactivation gate *q* was described by *q*_*inf*_(*v*) = {1 + exp [(*v* + 78.5)/6]}^−1^ and *r*_*q*_(*v*) = *k* * (*v* + 105). The conductance *gK_v_1* was decreased from 10 mS/cm^2^ to 5 mS/cm^2^, *k* was changed from 8.96 to 6.72. The surface area was increased by a factor of 25.2 and leak conductance per unit area increased by a factor of 2.5 to better account for the calculated capacitance and input resistance of the neurons and to better replicate the recorded f/I curve. Simulations were run in Neuron (Hines and Carnevale 1997), code is available at (https://modeldb.science/2016658), password is persistent_PV.

|  | Chamberland Model | Modification |
| --- | --- | --- |
| Length of cell | 10 μm | 126 μm |
| $g_L$ | 0.1 mS/cm<sup>2</sup> | 0.25 mS/cm<sup>2</sup> |
| $g_{Kv1}$ | 10 mS/cm<sup>2</sup> | 5 mS/cm<sup>2</sup> |
| Half activation of the $I_{Na}$ activation gate | -24 mV | -22 mV |
| *slope*-factor for $K_v1$'s inactivation gate | 10 | 7.5 |

To better capture the frequency-current relationship of PV+ fast spiking basket cells, the half activation of the sodium channel was changed from -24 mV to -22 mV. The conductance of the K_v_1 channel was decreased by a factor of 0.5 and the time constant of inactivation was decreased by 25%.

The model for the medial entorhinal cortical interneuron was taken from the homogeneous network in (Via et al. 2022) without modification. The model for the connected CA1 pyramidal cell was taken from (Bezaire et al. 2016) without modification. The PV+ cell synapsed onto the soma of the pyramidal cell and was described with a biexponential conductance with a rise time constants of 0.3 and 6.2 ms a total weight of 0.16764 and a conduction delay of 1 ms.

### Bifurcation Analyses

Bifurcation analyses were performed using by the MATCONT package (Dhooge et al. 2003).

## Results

### Model Calibration

To better capture the passive properties and frequency current relationship of hippocam-pal CA1 PV-INs, the model in (Chamberland et al. 2023b) was modified as described in the Methods. The resulting input resistance of the model was 80.6 MΩ, similar to that measured experimentally *in vitro* 78.7 MΩ (S.D.=24.4, n=27). The model had a membrane time constant of 4.1 ms and a capacitance of 50.9 pF, which were at the low end the experimental values (10.4 ± 4.9ms, n=27 and 142 pF ± 82.3, n=27) but still physiologically plausible. Depolarizing steps of increasing amplitude revealed a non-linear relationship between firing frequency and current injection in the model, closely aligned with experimental observations (Fig. 1A-C). The progression from non-spiking or transient firing to sustained high frequency firing resulting in discontinuous F-I curves in PV-INs is consistent with type 2 excitability as originally described by Hodgkin (Hodgkin 1948), a feature which our model is the first to capture in CA1 PV-INs.

**Fig. 1.**
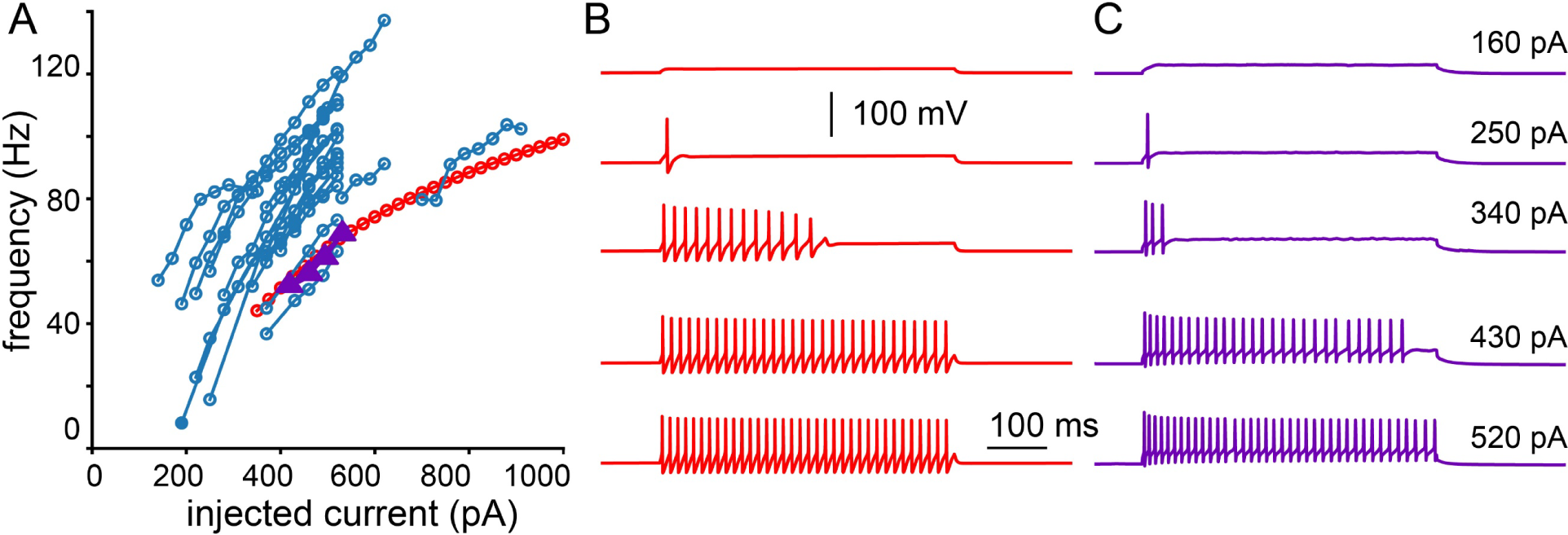
Frequency Current (f/I) Relationship. A. Experimental data points (blue circles, purple triangles). Model f/I curve (red circles) B. Voltage traces in model neuron in response to step currents. C. Voltage traces from experimental data points from a representative interneuron marked by purple triangles in A. At values of depolarizing current too low to support repetitive spiking, both model and real neuron emit one or more spikes then fall silent as inactivation is removed from K_V_1.

### Bifurcation analysis of Persistent Interruption

PV-INs firing can be persistently interrupted by brief membrane hyperpolarization, leaving the neuron in a depolarized quiescent state. We next aimed to obtain a mathematical understanding of the transitions between firing and silent states in PV-INs. The mechanisms underlying transitions between spiking and quiescence can be understood using a bifurcation diagram that graphs the transition points in a space consisting of a fast variable on the y-axis, and a slow variable on the xaxis (Izhikevich 2007).

Our model replicated the interruption (Fig. 2A top) using a simulated current clamp waveform (Fig. 2A bottom). The model neuron is initially silent but a 450 pA current step induces tonic firing (Fig. 1B) that eventually reaches a steady state (Fig. 2A, blue). During tonic firing, the available fraction of the K_v_1 channel (q) gradually accumulates until it reaches the steady state where the spike after-hyperpolarization (AHP) removes the exact amount of inactivation that occurs during the depolarized part of the action potential. At this point the value of q in Fig. 2B levels out, with a small residual oscillation around a steady value. The applied current was reduced sharply to 350 pA and linearly increased back to 450 pA during a 200 ms ramp to simulate an IPSP (a mock IPSP(Chamberland et al. 2023b). The neuron was silenced during the mock IPSP (red trajectory in Fig. 2A) but also remained silent for hundreds of milliseconds after the end of the mock IPSP (gray curve). The slow ramp was essential for the interruption to be observed. Subthreshold oscillations preceded the resumption in firing in experiments (Chamberland et al. 2023b Fig. 5AD and Supplementary Fig. 7E) and in the model, suggesting the emergence of a subcritical Hopf bifurcation (Izhikevich 2007).

**Fig. 2.**
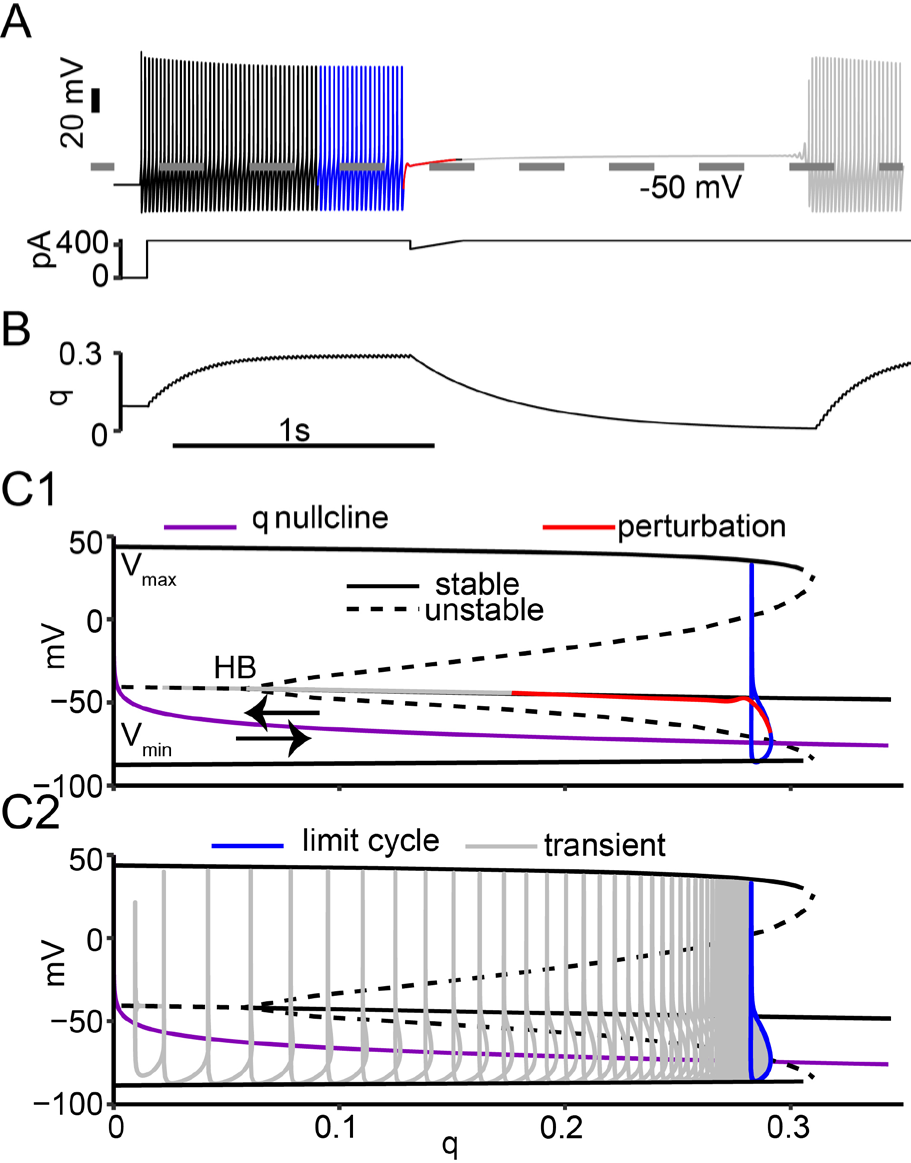
Bifurcation Analysis of Persistent Interruption. A. Persistent interruption of tonic firing by a mock IPSP in a model PV+ cell. B. Time course of K_V_1 inactivation gate. C. Bifurcation diagram with q as bifurcation parameter. C1. Leftward movement of trajectory during perturbation and interruption. C2. Rightward movement of trajectory during transient tonic spiking.

To better understand the firing interruption and firing resumption transitions in the system, we performed a bifurcation analysis (Fig. 2C) using fast/slow time scale analysis (Bertram and Rubin 2017). Since the inactivation gate for K_v_1 (*q*) builds up slowly during tonic firing and decays slowly during the firing interruption, we used *q* as a parameter in the bifurcation analysis. The bifurcation analysis holds the slow parameter constant to find the points at which transitions occur. The output of the bifurcation analysis was a set of fixed points, (resting membrane potentials) which could be stable (solid black curves) or unstable (dashed black curves), and the maxima and minima of oscillations in membrane potential (limit cycles). The subcritical Hopf bifurcation (HB in Fig. 2C1) at the transition point between the branches of fixed points. The q nullcline (purple curve in Fig. 2C), which is the steady value of q at each value of the membrane potential separates the regions of state space where q is increasing (below) and decreasing (above). These results show the dual role of q in establishing a stable firing state and an unstable dynamic state at the subcritical Hopf bifurcation.

Next, we consider *q* as dynamic variable to understand how the bifurcation structure drives the model trajectory. We superimposed the trajectory of the model from Fig. 2A and B into the *q* /V plane in Figs. 2C1 and C2, discarding the initial firing (black trace in Fig. 2A) and beginning at the steady state oscillation shown in blue in Fig. 2A and in 2C. A major modification in the Chamberland et al. 2023 model compared to the Golomb et al. 2007 model is in the description of K_V_1. In Golomb et al. 2007 the inactivation time constant (τ_q_) for K_V_1 is 150 ms at all membrane potentials. In Chamberland et al 2023 and in this study, the equation for τ_q_ depends linearly on membrane potential, so it is large (slow) at the depolarized potentials at which the membrane spends most of its time, but faster during the AHP at hyperpolarized potentials. The q nullcline divides the q /V plane into two halves. Below the nullcline, in the simulations q increases rapidly and the trajectory moves to the right. Above the *q* nullcline, *q* slowly decreases moving the trajectory to the left. This dynamic picture allows us to make sense of the model activity, as explained below.

At the subcritical Hopf bifurcation (Izhikevich 2007), fixed point (resting membrane potentials) transitions from unstable (dashed line) to stable (solid curve obscured by the red and gray trajectory curves in Fig. 2C1 but visible in Fig. 2C2). In the bifurcation diagram, an oscillation is described only by the maximum and minimum values of membrane potential achieved. An oscillation that traces an identical cycle on each oscillation is called a limit cycle. An unstable limit cycle (indicated by the two dashed curves representing the maxima and minima) which is not robust to noise (Izhikevich 2007) emerges from the subcritical Hopf bifurcation (HB). The unstable limit cycle becomes a stable oscillation representing tonic firing (maxima and minima indicted by the two solid curves) and forms the boundary between the stable repetitive firing and a stable resting membrane potential that coexist at values of *q* held constant between about 0.07 and 0.3. This coexistence is called bistability. A neuron that starts its trajectory outside of this unstable limit cycle will continue firing on the stable limit cycle, but trajectories within the unstable limit cycle move towards the fixed point. Since the model is bistable, the interruption (shown in red) pulls the model’s trajectory within the unstable limit cycle and lands on the branch of stable fixed points representing the depolarized quiescent state. In repetitive spiking, the fast Na^+^ current produces an action potential upstroke before the K^+^ currents can activate sufficiently (and the Na^+^ channel inactivates sufficiently) to prevent the action potential. Although the neuron is depolarized beyond the action potential threshold observed during repetitive firing, it remains silent because the slow return to the baseline value of injected current allowed the fast variables to reach a steady state in which the outward currents dominate. The trajectory then moves slowly to the left through a series of stable fixed points (that are only stable at a constant value of *q*) as *q* decays. This series of “stable” fixed points was referred to by Chamberland et al. 2023 as a “drifting stable point in the membrane potential”; here we show that the drift is caused by the slow dynamic evolution of the *q* variable, resulting in an interruption that outlasts the mock IPSP. After *q* decays below the Hopf, the trajectory travels along the weakly repelling branch of unstable fixed points, which prolongs the interruption, until the repulsion becomes obvious as an oscillation that starts with small amplitude (inset in Fig. 2C1) and grows. Fig. 2C2 illustrates the trajectory once action potential firing commences, recapitulating the transient activity (gray trace) as *q* increases moving the trajectory to the right, until steady repetitive firing (limit cycle shown in blue) is achieved.

### Elliptical Bursting

Elliptical bursting (Rinzel and Ermentrout 1998; Wang and Rinzel 1995), originally called type 3 bursting (Bertram et al. 1995; Rinzel 1987), requires a slow variable to drive the intrinsic dynamics back and forth across a subcritical Hopf bifurcation. PV-INs can undergo firing states reminiscent of elliptical bursting, sometimes referred as to ‘’stuttering’’ (Bracci et al. 2003; Chamberland et al. 2023b; Markram et al. 2004; Sciamanna and Wilson 2011). We next explored if transitions between firing and silent states occur similarly as to the firing interruption.

The model exhibited elliptical bursting in response to moderate current injection (Fig. 3B), consistent with the observation of noisy elliptic putative bursting in experiments (Fig. 3B1). The availability of K_V_1 indicated by the gating variable *q* (Fig. 3B2) controls the timing of the bursts. Fig. 3B2 shows the bifurcation diagram with q held constant at an injected current of 330 pA. As in Figure 2C (calculated at 450 pA injected current), the inactivation gating variable *q* accumulates rapidly during the AHP that drops below that nullcline, then decays slowly during the subsequent depolarization and action potential. However unlike in Figure 2C, a tonic, steady firing rate is never achieved. Instead, during spiking *q* continues to increase until the branch of stable limit cycles (solid curves representing the maxima and minima of the membrane potential) collides with the branch of unstable limit cycles at a bifurcation called a saddle node of periodics (Izhikevich 2000). Since no stable limit cycle exists above a *q* value of about 0.18 (for this level of current injection), the trajectory collapses onto the branch of stable fixed points (not visible under the blue trajectory but similar to that seen in Fig. 2C2). The trajectory then slowly relaxes to the left as the fixed point is above the *q* nullcline along a trajectory of stable fixed points (with respect to a fixed *q*) as *q* decays. When *q* decays past the Hopf, the stable membrane resting potential becomes unstable, initiating another cycle of spiking (Knowlton et al. 2020). These results show that PV-INs are silenced by distinct mechanisms during the firing interruption and elliptical bursting, while the return to the firing state is similarly explained in both cases by a drift away from the Hopf driven by the slow dynamic evolution of *q*.

**Fig. 3.**
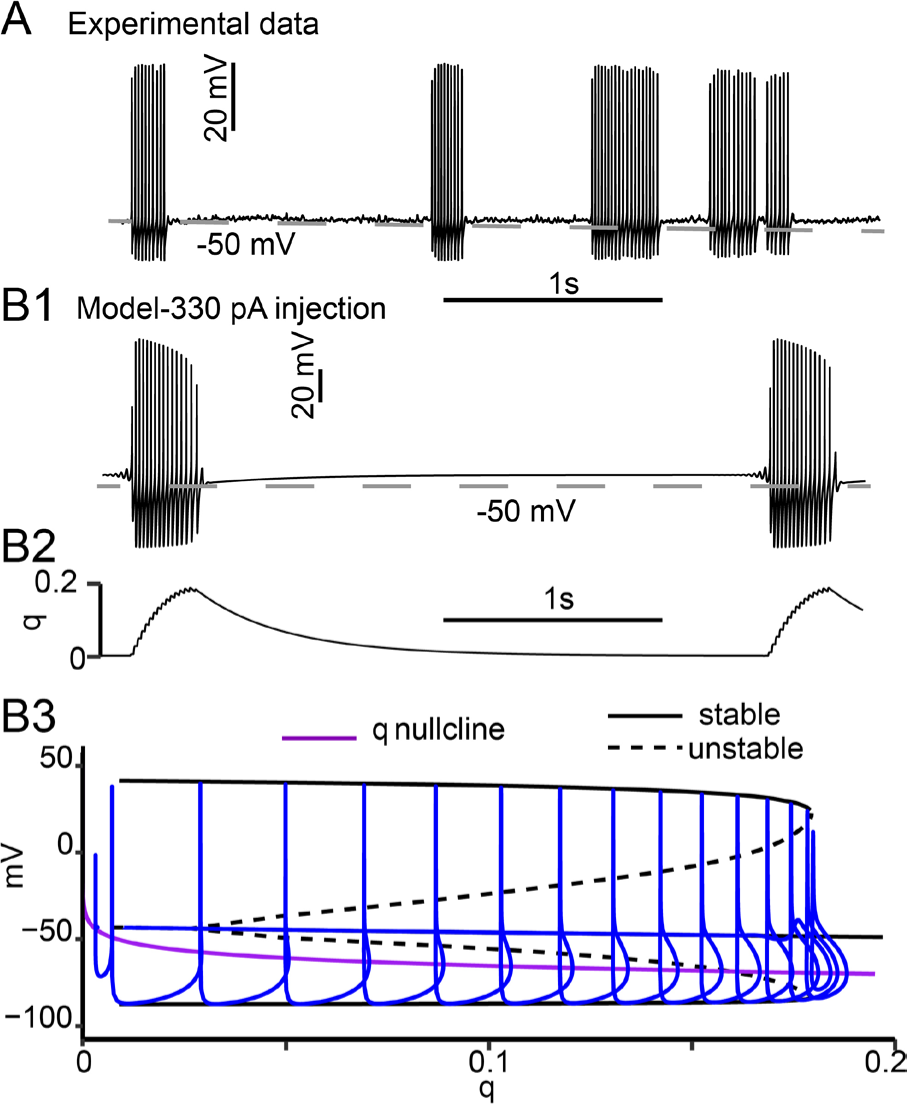
Elliptical Bursting. A. Experimentally observed trace recorded in BIC + CGP. Experimentally observed elliptic bursting is less regular than that of a noiseless model because of the physiologically unavoidable noise although these experiments were performed in the presence of blockers of inhibitory synaptic transmission. B1. Model membrane potential showing repetitive elliptical bursting with same parameters as Fig. 1C and 2 except the injected current was 330 pA. B2. Time course of *q*. B3. Model trajectory for a single burst superimposed on bifurcation diagram.

### Parameter Space Compared to Neocortical PV-IN Model

The above results implicate the existence of different firing regimes in CA1 PV-INs, as previously described in a model of neocortical PV-INs (Golomb et al, 2007). Given that this model was based on Golomb et al. (2007), we next explored the model to obtain a description of the different firing regimes in CA1 PV-INs.

We first explored the parameter space in the plane of gK_V_1 and Iapp. We varied the injected current from 0 pA to 1000 pA in 30 pA increments, and the conductance of K_v_1 from 0 to 10 mS/cm^2^ in increments of 1 mS/cm^2^. Similar to Fig. 2 of Golomb (2007), we observed four different firing regimes (Fig. 4A). Either the model was quiescent, fired transiently, exhibited elliptical bursting or fired continuously. Figure 4B shows an f/I curve with gK_V_1 set to zero (bottom), and an example of tonic firing (top). Since the f/I curve is discontinuous with an abrupt onset of firing at about 40 Hz, in our model K_V_1 is not required for type 2 excitability. Figure 4C gives an example of a transient response. In contrast to Golomb (2007), transient spiking prior to quiescence was always observed instead of a delay prior to initiation of tonic spiking. This difference is explained by the description of K_V_1. Since there is some overlap in the steady state activation and inactivation curves for K_V_1, there is a small steady state “window” current at membrane potentials near the resting membrane potential. Here, K_V_1 has a much smaller window current and therefore a much smaller fraction of available K_v_1 at rest. In the Golomb (2007) model, there is sufficient K_V_1 available at rest to silence the cell for sufficiently small values of Iapp, hence explaining the delay. Firing can only commence after inactivation of the K_v_1 channel at the more depolarized potential resulting from the injected current. With less K_v_1 available at rest, there is insufficient K_v_1 current to stop the model from firing with the initial current injection. During the initial bout of spiking, the inactivation is removed during the AHP, allowing outward K_V_1 current to accumulate and render the model quiescent. This finding reconciles observations that PV-INs can exhibit delayed firing or fire at the onset of the depolarizing pulse. Considering that K_v_1 properties can be activity-dependent, this would provide a likely explanation for differences between PV-INs. Figure 4D gives an example of elliptic bursting. Elliptical bursting in both models required a minimum amount of gK_v_1. In contrast to Golomb et al. 2007, elliptic bursting began in the spiking phase rather than the quiescent phase. The difference in window current at rest also accounts for the lack of delay prior to initiating spiking during elliptic bursting in our model. Despite this difference, our simulations showed that that the Golomb model in Fig 2C from that paper could also exhibit persistent interruption (data not shown).

**Fig. 4.**
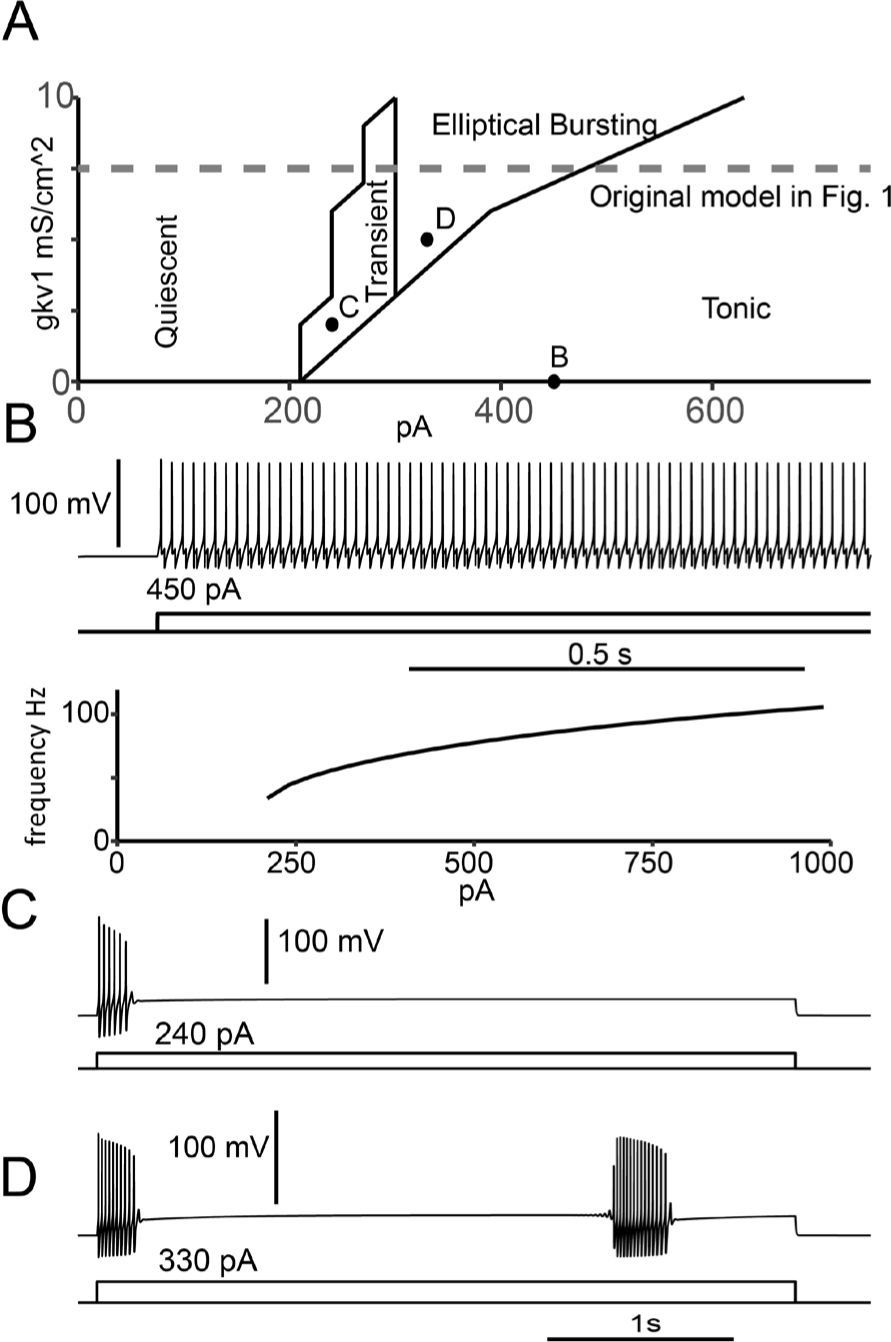
Parameter Space. A. Phase diagram in the plane of g_K_V_1_ and I_app_. B. Top: tonic firing at 0 mS/cm^2^ and 450 pA Bottom: Discontinuous f/I curve at 0 mS/cm^2^ characteristic of type 2 excitability (see text). C. Transient firing at 2 mS/cm^2^ and 240 pA. D. Elliptic bursting at 5 mS/cm^2^ and 330 pA repeated from Fig. 3B1.

### Hopf bifurcation parameter: applied current versus slow dynamic variable

To understand how widespread the interruption of firing is across PV-INs models, we challenged a previously published model from the medial entorhinal cortex with similar simulation paradigms (Via et al. 2022). Both models exhibit type 2 excitability, identified by the subcritical Hopf bifurcation with Iapp as the bifurcation parameter; this bifurcation occurs at the value of Iapp at which tonic spiking is initiated as Iapp is increased. This bifurcation is unrelated to the subcritical Hopf bifurcation in our model of a CA1 PV-IN that occurs considering *q* as the bifurcation parameter. The subcritical Hopf bifurcation that underlies type 2 excitability implies a region of bistability between spiking and quiescence at values of Iapp on the stable side of the bifurcation.

At certain values of Iapp and with an appropriately calibrated mock IPSP, a persistent interruption of firing can also be demonstrated in the (Via et al. 2022) model (Fig 5A1 and B1). Annihilation of firing that lasts beyond stimulus termination was first demonstrated in squid axon (Guttman et al. 1980), the very preparation in which Hodgkin and Huxley first modeled the generation of action potentials. Their classic model (Hodgkin and Huxley 1952) exhibited type 2 excitability due to a subcritical Hopf bifurcation, and this bistability was invoked to explain the observed cessation of firing in response to an appropriate perturbation. Therefore, both models of PV-INs could demonstrate the interruption of firing.

**Fig. 5.**
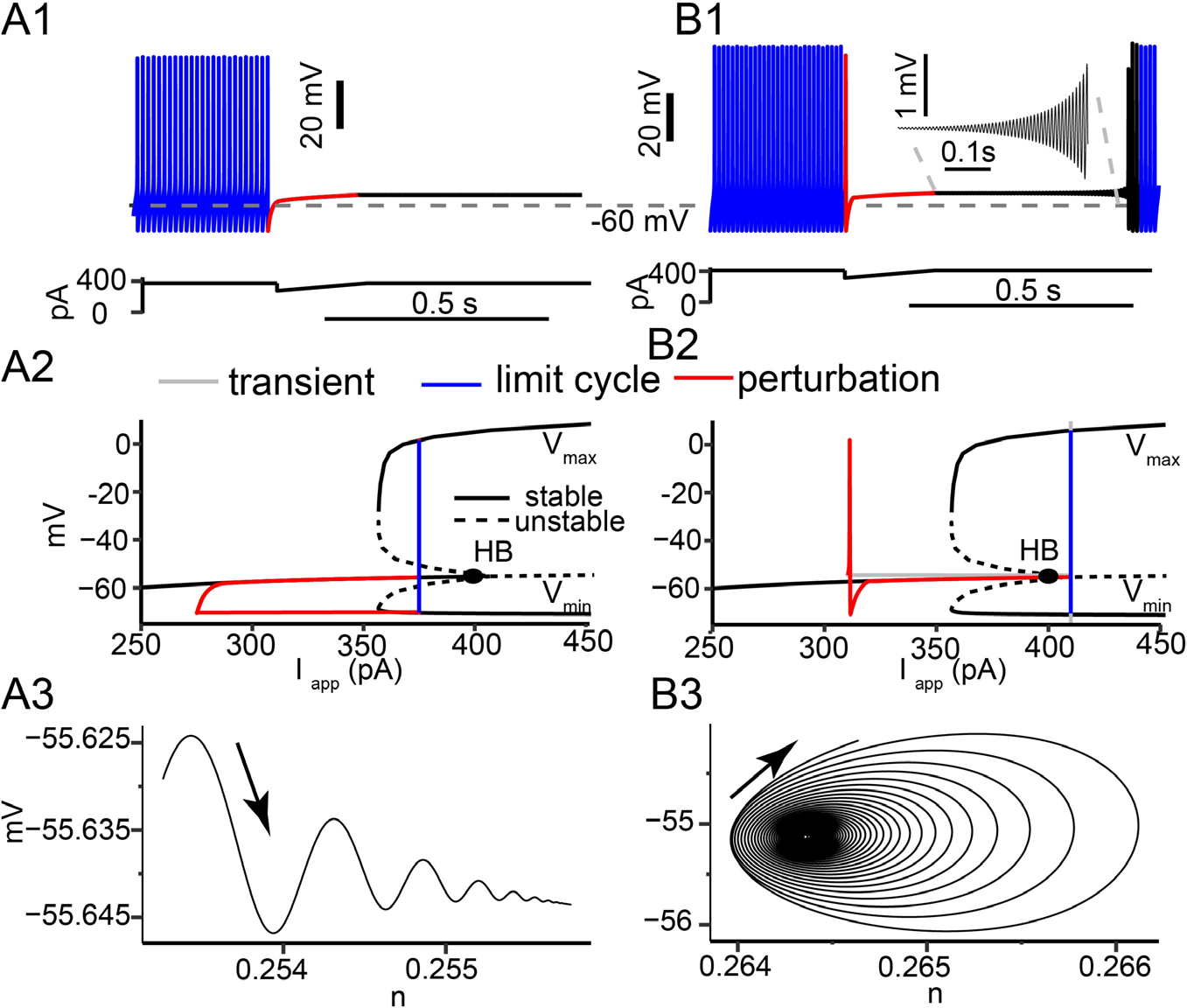
Perturbation from tonic firing to a stable versus an unstable branch of fixed points. The Via model at two levels of current, within the bistable regime (**A**) and a globally attracting tonic firing regime (**B**). A1. A mock IPSP of maximum amplitide100 pA that decayed linearly within 200 ms was subtracted from a constant applied current of 375 pA. A2. The trajectory was superimposed on the bifurcation structure of the model with applied current as the slow parameter. A3. Damped oscillatory trajectory in the phase-plane of the n gate for Kv3 and the voltage as the stable fixed point is approached during the transient following the interruption. B1. A similar mock IPSP was applied at a holding current of 410 pA. B2. Trajectory superimposed on the bifurcation structure. B3. Trajectory in the phase plane of the n gate for Kv3 and voltage spiraling out from the branch of weakly repelling fixed points during the transient following the interruption.

We next investigated the mechanisms controlling the interruption of firing in both models. In the EC PV-IN model at a holding current of 375 pA, a mock IPSP produces a permanent transition from tonic firing to quiescence in a noiseless model (Fig. 5A1). The bifurcation diagram in Fig. 5A2 has Iapp, the applied current, as the bifurcation parameter instead of the gating variable q as in Fig. 2C and 3B3. Fig. 5A2 shows that at a holding current of 375 pA, the limit cycle representing tonic firing is just to the left of the subcritical Hopf; therefore, tonic firing is bistable with a stable resting membrane potential. The perturbation by the mock IPSP (red) moves the trajectory onto the stable branch of fixed points. Here, since Iapp is a parameter that is held constant except during the perturbation, the fixed points are truly stable. Thus, in the absence of noise the trajectory remains on the stable branch after the perturbation, with just a brief small amplitude transient (Fig. 5A3). In contrast, at an injected current of 410 pA, a mock IPSP elicits a persistent but finite interruption of firing (Fig. 5B1). The finite duration interruption is a consequence of the increased injected current shifting the limit cycle trajectory to the right of the Hopf bifurcation (Fig 5B2). Thus, the model is no longer bistable. The hyperpolarizing input (red) again moved the trajectory onto the branch of stable fixed points. However, once the trajectory crosses the Hopf bifurcation as the applied current returns to its original depolarized value, the trajectory is forced to spiral out (Fig. 5B3) into the limit cycle that represents tonic repetitive firing. Although the branch of fixed points is unstable beyond the Hopf bifurcation, it is only weakly repelling due to the small amplitude of the positive real parts of the eigenvalues at the fixed points (Izhikevich 2007). The spiral is very slowly expanding (inset in Fig. 5B1), resulting in the persistent interruption.

Our results suggest that the main effect of these differences would result in increased resilience to noise. To inject a comparable amount of noise in each model, we set the standard deviation of zero mean injected Gaussian current noise to the amount of current required to hyperpolarize each model by 5 mV (Destexhe and Rudolph-Lilith 2012) from its Hopf bifurcation point, assuming that *q* is held at its equilibrium value at the Hopf bifurcation (0.05764) in the Chamberland model. This noise was added to the applied current waveform to compare the sensitivity of the different types of interruption to noise. In each case, ten different noisy simulations were run. Fig. 6A2 shows that the duration of the interruption in the Chamberland model from Fig. 2A is still several hundreds of milliseconds (mean interruption length 555.2 ms with standard deviation of 39.5 ms) in the presence of substantial current noise, although only about half the duration in a noiseless simulation (Fig. 6A1). In contrast, in the Via model in Fig. 6B1 (repeated from Fig. 5B), which was not biased in the bistable range, the interruption vanishes (3.4 ms with standard deviation 16.9 ms) in the presence of substantial current noise Fig. 6B2). The trajectory during the mock IPSP is shown in red, and in this example firing resumes slightly before the mock IPSP terminates. In the Via model in Fig. 6C1 (repeated from Fig. 5A), which was biased in the bistable regime, the interruption was also not robust to noise, but in a different way. In all ten noisy simulations, including the one shown in Fig. 6C2, an interruption was triggered by noise before the mock IPSP was delivered, and in 8 of 10 there were multiple transitions between spiking and quiescence. Therefore, we found that the mechanisms controlling the interruption of firing were not the same in the hippocampal and EC PV-IN models.

**Fig. 6.**
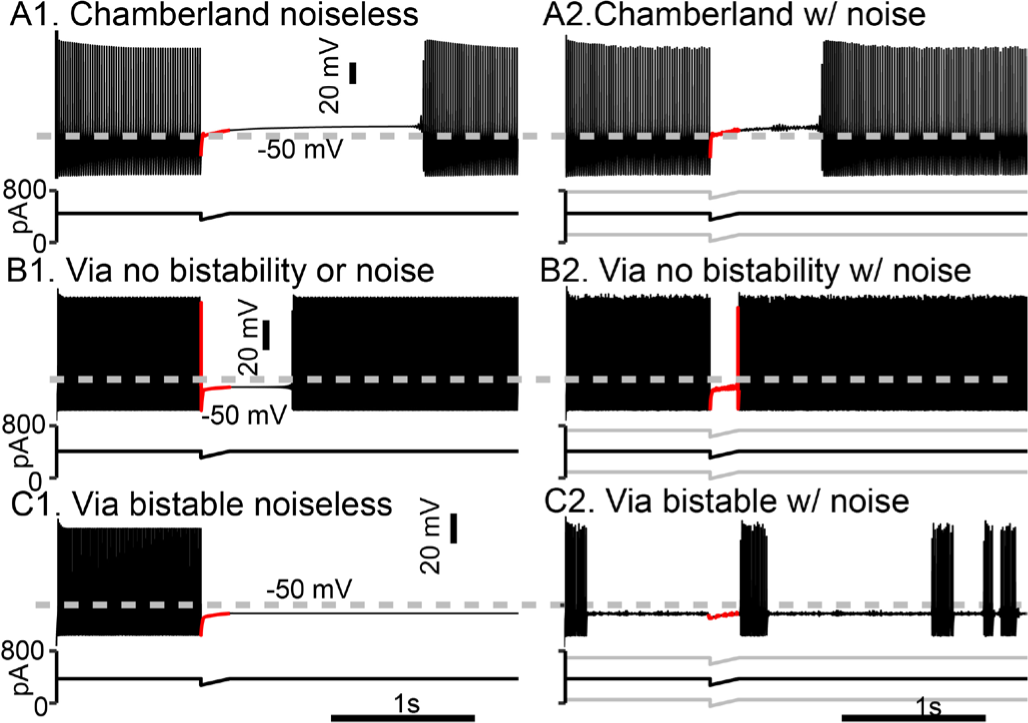
Robustness of interruptions to noise. The Chamberland model was compared to the Via model at two levels of current. The mock IPSP was 100 pA over 200 ms in all cases. The voltage waveform during the mock IPSP is shown in red. A1. Persistent interruption of noiseless Chamberland model repeated from Fig. 2A. A2. Same as A1 but added Gaussian current noise with a mean of zero and a standard deviation of 164.7 pA. B1. Persistent interruption in noiseless Via model repeated from Fig. 5B1. B2. Same as B1 but with additive Gaussian current noise with a standard deviation of 160.6 pA. C1. Persistent interruption in Via model repeated from Fig. 5A. C2. Same as B1 but with additive Gaussian current noise with a standard deviation of 160.6 pA. Switches between firing and quiescence occur independently of the mock IPSP.

The level of noise or synaptic input required to cause the Chamberland model to return to firing is proportional to the size of the unstable limit cycle. The bifurcation diagrams in Fig 2C and Fig 3B3 shows that in the bistable regions, the unstable limit cycle forms the boundary between quiescence and tonic firing. In Fig 7A (repeated from Fig. 6A1 with the same parameter settings as Fig. 2A), the mock IPSP pulls the trajectory within the unstable limit cycle, and the model becomes quiescent. During this interruption, an additional EPSC (excitatory postsynaptic conductance) can push the trajectory back outside the unstable limit cycle, inducing firing as demonstrated in Fig 7B. The same EPSC is unable to induce firing in the model at rest (Fig. 7C). As *q* recovers, the unstable limit cycle shrinks and the minimum peak conductance EPSC to induce firing decreases (Fig 7D). Because the model is bistable, the required stimulus to induce firing is generally less than required at rest. The unstable limit cycle is a curve that surrounds the fixed points of the model both in the depolarizing and hyperpolarizing directions. Therefore, an inhibitory stimulus that pushes the trajectory past the hyperpolarized boundary of the unstable limit cycle will also induce firing. The required minimum conductance of this stimulus decreases during the interruption (Fig. 7E), due to the shrinking of the unstable limit cycle as *q* recovers.

**Fig. 7.**
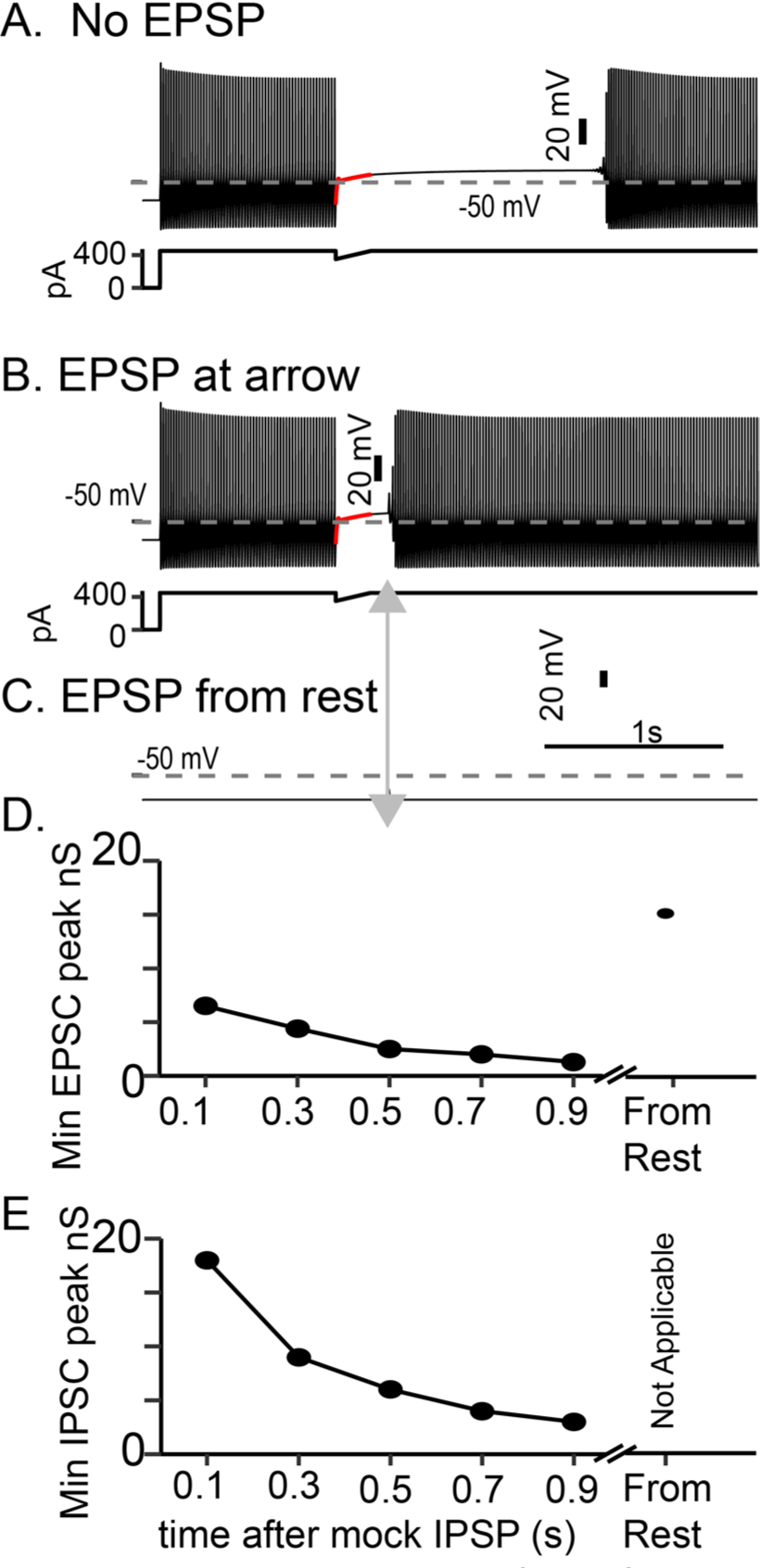
The quiescent state of the Chamberland model becomes more susceptible to synaptic input over time. The Chamberland model, when interrupted from rapid firing by an inhibitory mock IPSP enters a quiescent state (**A**). Excitatory synaptic input given shortly after the end of the mock IPSP can return the model to firing (**B**). In B, a biexponential conductance with a rise time of 0.5 ms, a fall time of 1 ms, a reversal potential of 0 mV and a maximum conductance of 6.4 nS was enough to cause the model to return to firing. The same synapse given to the model at rest (**C**) did not. The maximum required conductance for the model to return to fire decreases during the interruption (**D**), and is higher at rest. As the model is bistable in the quiescent state, an inhibitory synapse with a reversal potential of -65mV can also cause the model to fire, though it is not sufficient at rest. The maximum required conductance for an inhibitory synapse also decreases with the duration of the interruption (**E**).

### Network Implications

Since PV-INs target the perisomatic region of CA1-PYRs, the persistent interruption could lead to an increase in firing locally connected pyramidal cells, as shown experimentally by (Chamberland et al. 2023b). We replicated this phenomenon, by connecting our model to a pyramidal cell model taken from (Bezaire et al. 2016), with inhibitory synapses targeting the soma. Similar to (Chamberland et al. 2023b), we injected enough current into the pyramidal cell to elicit tonic firing, and a baseline current and mock IPSP waveform into the presynaptic PV-IN cell. The locally connected pyramidal cell fires tonically while the PV-IN cell is quiescent, falls silent while the PV-IN cell is firing, and resumes firing when the PV-IN cell is interrupted (Figure 8). This leads to a persistent increase in pyramidal cell firing, during the persistent interruption in PV-IN. We suggest this could be a mechanism for short-term memory, in which a transient stimulus causes a persistent increase in activity which is constrained here by the duration of interneuronal silence.

**Figure 8.**
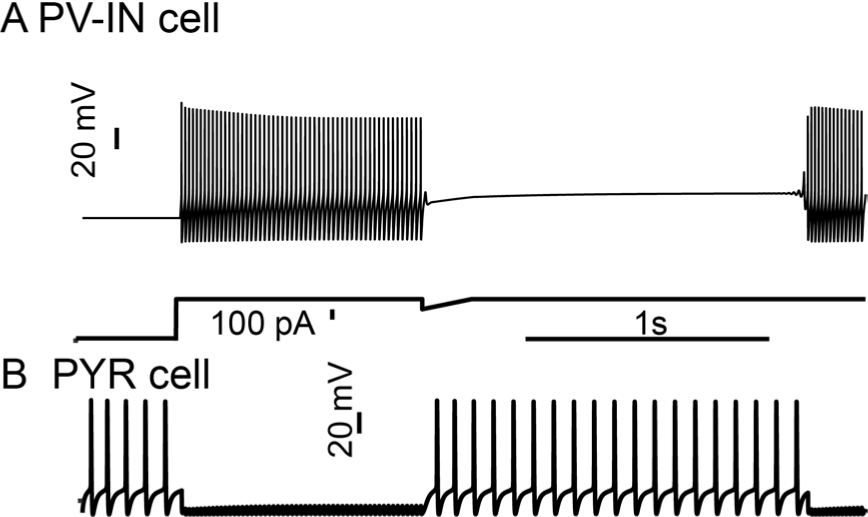
Local network implications. A. The PV-IN model (top) was injected with 450 pA and a 100 pA mock IPSP was applied (bottom). B The pyramidal cell with 275 pA applied current to elicit tonic firing. The PV-IN model synapsed on the PYR model, such that when the PV-IN cell was active, it silenced the PYR model.

## Discussion

### Summary

Persistent interruption of firing in PV-INs was recently demonstrated (Chamberland et al. 2023b). In this study, we used a fast/slow separation of times scales approach with the K_v_1 inactivation variable as the bifurcation parameter to show that an inhibitory post-synaptic potential can stop repetitive firing by moving the trajectory onto a co-existing stable fixed point corresponding to a depolarized quiescent state. Although the interruption could be observed in all models of PV-INs tested, the slow inactivation of K_v_1 at depolarized potentials made persistent interruptions resistant to noise and E_Na_bled elliptical bursting at some levels of depolarization. The persistent interruption of PV-INs firing can potentially trigger persistent activity in post-synaptic pyramidal cells via disinhibition.

### FS neurons display Type 2 excitability

Early models of CA1 fast-spiking PV-INs (Wang and Buzsáki 1996) as well as more recent ones (Ferguson et al. 2013) exhibited Hodgkin’s type 1 excitability, characterized by an f/I curve that theoretically starts at arbitrarily low frequencies (Hodgkin 1948; Izhikevich 2007; Knowlton et al. 2020). On the other hand, models of fast spiking neurons in mouse somatosensory cortex (Erisir et al. 1999), like their experimental counterparts, exhibited type 2 excitability with a minimum frequency of tens of Hz. Type 2 excitability provides a mechanism for “stuttering” that is distinct from elliptic bursting, although in the presence of noise they appear similar. In the bistable region (Fig. 6C2), noise can induce switches between quiescence and tonic firing (Izhikevich 2007; Schreiber et al. 2009). Bistability based solely on type 2 excitability is not a reliable substrate for persistent interruption, as shown in Fig. 6. An extension of the Erisir model (Golomb et al. 2007) examined different parameter settings regarding the contribution of the Na^+^ window current that controls whether inward or outward currents dominate at steady state in the range of membrane potentials traversed during the interspike interval. Dominant inward currents produce type 1 excitability and dominant outward currents produce type 2 excitability (Prescott et al. 2008). It is possible that PV-INs exhibit both types of excitability depending upon their exact mix of conductances, but our data suggest predominantly type 2 excitability in mEC (Martínez et al. 2017; Tikidji-Hamburyan et al. 2015; Via et al. 2022) and area CA1 (Fig. 1).

### Generalized Role of K_V_1 in Persistent Interruption and Transient Firing

K_v_1 channel blockers eliminated the delay to first spike observed at values of Iapp near the threshold to evoke tonic firing in layer 2/3 mouse barrel cortex (Goldberg et al. 2008). These delays result from inactivation of K_V_1 channels and are commonly observed in neocortical PV-INs (Erisir et al. 1999; Golomb et al. 2007) but not in medial entorhinal cortex (mEC) (Via et al. 2022) or CA1 PV-INs (Fig. 1BC and Chamberland et al. 2023), and a mixed phenotype appears to be observed in striatal PV-INs ((Bracci et al. 2003). Modeling suggests that a smaller window current at rest eliminates the delays by making less K_V_1 conductance available. In the (Golomb et al. 2007) model after a current step, it takes time for K_v_1 to inactivate and therefore there is a delay in firing. In our model there is less K_v_1 at rest, so our model starts to fire immediately during a current step, but for sufficiently small current steps, enough inactivation is removed from K_v_1 during the AHPs to silence the neuron. Therefore instead of delays at values of Iapp near the threshold for tonic firing, in CA1 (Fig. 1BC and Chamberland et al. 2023) and mEC (Via et al. 2022), transient firing is observed in which one or more spikes are emitted before the neuron falls silent. In CA1, the model therefore predicts that K_V_1 blockers would convert the transient firing to tonic firing as in (Sciamanna and Wilson 2011). Our model and the experimental observations in area CA1 broaden the repertoire of activity conferred on PV-INs by K_V_1 expression.

The mechanisms for transitions between firing and quiescence (Chamberland et al. 2023b) are similar to those observed previously (Golomb et al. 2007; Sciamanna and Wilson 2011) in models that also include the slowly inactivating K_V_1 current, but distinct from those in another model (Via et al. 2022). The Via model exhibits a less robust persistent interruption of firing due to the subcritical Hopf bifurcation underlying type 2 excitability, which does not involve K_V_1 current whose slow time scale protects the interruption from termination by noise. Therefore, K_V_1 not only E_Na_bles persistent interruption, it also contributes to type 2 excitability and may in some cases be required, as in striatal PV-INs (Sciamanna and Wilson 2011).

In the absence of additional information, Via et al. 2022 modeled the transient firing as terminating due to slow activation of a K^+^ current they called K_V_1, but whose characteristics more closely resemble the slowly-activating K_V_7 (Prescott and Sejnowski 2008). Experiments using K_V_1 and K_V_7 blockers are required to determine which current is responsible for cessation of firing of PV-INs in the mEC, but it seems likely that the mEC interneurons, like their cortical and hippocampal counterparts, rely on K_V_1 rather than K_V_7 to regulate firing activity near threshold. Thus, we predict that robust persistent interruption of firing can be observed in mEC and neocortical PV-INs, just as in CA1. Moreover, K_v_1 channels are expressed in many other neurons including dopaminergic neurons (Fulton et al. 2011), and deep cerebellar neurons (McKay et al. 2005). Therefore, it is possible that other neuronal types may exhibit some of the features E_Na_bled by K_V_1 such as transient firing, firing delays, elliptic bursting and persistent interruption of firing.

### Inhibitory dynamics controlling CA1 activity

CA1-PYRs often function as place cells that fire when an animal is in a particular area of their environment (O’Keefe and Dostrovsky 1971), and have been suggested as model for general episodic memory (Redish 1999). Local CA1 inhibitory circuits contribute to place cell firing (Grienberger et al. 2017; Royer et al. 2012; Valero et al. 2022; Zutshi et al. 2022),and interneurons are thought to coordinate sequences of place cells (Udakis et al. 2020; Valero et al. 2022; Zutshi et al. 2022). The influence of different subtypes of interneurons on place cells firing varies as a function of place field position. This is best exemplified by the progressive shift in the influence of PV->SST INs over CA1-PYR firing when the animal traverses a place field (Royer et al. 2012), for which PV-mediated inhibitory influence on CA1-PYR firing is highest at place field entry and gradually diminishes (Royer et al. 2012). This results in a progressive shift in perisomatic to dendritic inhibition, and it is remarkable that the CA1 circuit is intrinsically wired to support this shift (Pouille and Scanziani 2004).

The persistent interruption of firing is an attractive cellular mechanism to explain how the dampening of PV-mediated inhibition could enhance CA1-PYRs firing in their place field (Royer, 2012; Valero, 2022), likely complementing the short-term synaptic dynamics in the circuit (Pouille and Scanziani 2004). While any source of inhibition that is sufficiently strong can theoretically interrupt PV-IN firing, recent work (Chamberland et al. 2023a) identified a population of bistratified somatostatin positive (*Sst*) INs that expresses the *Tac1* (tachykinin precursor 1) gene. These *Sst;;Tac1*-INs preferentially synapses onto FS-INs and suffice to interrupt their firing, unlike two populations of *Sst*-expressing oriens lacunosum-moleculare (OLM) INs that predominantly target CA1-PYRs and for the most part avoided FS-INs. Thus, *Sst;;Tac1*-INs potentially control persistent interruption in PVINs and the resultant lasting disinhibition of CA1-PYRs. The stimulus precipitating persistent activity in this case would be the excitation of *Sst;;Tac1*-INs, known to fire after PVINs during hippocampal CA3 activity (Chamberland et al. 2023a). Increasing the depolarizing current injected into PV-INs abolished the ability of IPSPs to interrupt firing (Chamberland et al. 2023b). Therefore, it is likely that only the most weakly-excited PV-INs would be interrupted by *Sst;;Tac1*-INs, effectively disinhibiting subsets of CA1-PYRs. Disinhibition of CA1-PYRs subsets could also be achieved through variable interruption duration across PV-INs (Börgers et al. 2010) or by non-uniformly distributed synaptic inhibition strength provided by Sst;;Tac1-INs to PV-INs (Chamberland et al. 2023a). Thus, persistent interruption could be a substrate for a memory trace.

## Acknowledgments

This work was supported by NIH R01 NS054281 and NIH R01 MH115832 to CCC and NIH K99 MH126157 to SC.

